# A Computational Design of a Programmable Biological Processor

**DOI:** 10.1101/2020.03.04.976290

**Authors:** Miha Moškon, Žiga Pušnik, Lidija Magdevska, Nikolaj Zimic, Miha Mraz

## Abstract

Basic synthetic information processing structures, such as logic gates, oscillators and flip-flops, have already been implemented in living organisms. Current implementations of these structures are, however, hardly scalable and are yet to be extended to more complex processing structures that would constitute a biological computer.

Herein, we make a step forward towards the construction of a biological computer. We describe a model-based computational design of a biological processor, composed of an instruction memory containing a biological program, a program counter that is used to address this memory and a biological oscillator that triggers the execution of the next instruction in the memory. The described processor uses transcription and translation resources of the host cell to perform its operations and is able to sequentially execute a set of instructions written within the so-called instruction memory implemented with non-volatile DNA sequences. The addressing of the instruction memory is achieved with a biological implementation of the Johnson counter, which increases its state after an instruction is executed. We additionally describe the implementation of a biological compiler that compiles a sequence of human-readable instructions into ordinary differential equations-based models. These models can be used to simulate the dynamics of the proposed processor.

The proposed implementation presents the first programmable biological processor that exploits cellular resources to execute the specified instructions. We demonstrate the application of the proposed processor on a set of simple yet scalable biological programs. Biological descriptions of these programs can be written manually or can be generated automatically with the employment of the provided compiler.

## 1 Introduction

Synthetic biology aims to design biological information processing structures [1, 2], which advance the field with new applications and enhance the current bioengineering applications. These include biosensing and actuating, smart therapeutics and engineering for biofuels, drugs or biomaterials [3]. Moreover, synthetic information processing structures enable us to increase our understanding of cellular information processing [4]. These structures include logic gates [5], oscillators [6] and memory [7]. One of the goals in the field is to design these structures in a robust and reliable manner, which would enable us to implement synthetic information processing systems at least as scalable as simple digital electronic systems. These would enable us to implement a biological computer.

Transcription-based logic circuits, in which engineered DNA sequences presenting the transcription factor binding sites, promoters and transcription factor coding sequences are used to perform the computation, have been developed since the so-called *foundational years* of synthetic biology [8]. The first two synthetic constructs implemented *in vivo* in *Escherichia coli* in fact presented two structures with information processing capabilities, namely a *repressilator* [9] and a *toggle switch* [10]. These types of simple biological information processing structures have been adapted and extended in many different directions, e.g., to operate in mammalian cells [11, 12], to reflect more predictable behaviour [13], to work in a consortium of interacting cells [14, 15, 16, 17, 18], etc.

Even though the number of reported novel synthetic structures with information processing capabilities is still increasing, the complexity of these structures is far from their electronic counterparts. The main problem is in their limited scalability [1]. One of the main differences between the digital electronic systems and biological systems is in the free diffusion of signals carrying the information in the latter. It is thus vital to ensure orthogonality between the synthetic parts as well as orthogonality between the synthetic parts and the cellular metabolism of the host, which is extremely hard when the number of synthetic parts is increased [19, 20]. Several approaches, such as the use of synthetically designed transcription or translation regulators (see e.g. [21, 22]), engineered proteases [23] or a decomposition of a logic circuit among different cells in a population can be applied to solve this problem. Decomposition of transcriptional logic to less complex functions distributes the metabolic burden among different cells [2]. Moreover, distribution of the same functions among several cells, which are coupled through quorum-sensing or other cell-to-cell communication mechanisms, decreases the cell-to-cell variability and increases the robustness of the overall cellular response [24].

Recently, a lot of focus has been devoted to the implementation of logic structures using engineered proteases, which enable us to implement protein-based processing signalling logic [23]. The most recent advances include the construction of *split-protease-cleavable orthogonal-coiled coils-based* (SPOC) logic circuits [25] and *circuits of hacked orthogonal modular proteases* (CHOMP) [26]. These can be used to design artificial signalling functions, which operate independently of the slower cellular processes, such as transcription and translation, and thus have a much faster response time than transcription-based logic. These circuits have been used to implement Boolean logic functions [25, 26] as well as biomolecular switches [27]. However, their capacity to construct more complex circuits and circuits with periodic behaviour has not been investigated yet and might prove to be problematic due to irreversibility of proteolysis.

Digital electronic systems are able to achieve highly scalable complexity with their implementation as synchronous sequential circuits. In synchronous sequential circuits changes of the system are performed only at specific events. A *clock signal* is used to synchronize the system and switches from one state to another. Digital electronic systems are usually synchronized to the *positive edge* or the *negative edge* of the clock signal. This means that the state of the system cannot change more than once per clock period, enabling us to ignore certain drawbacks of digital electronics (such as signal propagation delays), and ensuring a reliable and predictable behaviour of the system [28].

Even though some theoretical work has already been reported in the field of synchronous sequential synthetic biological system (see e.g. [29, 30]) this concept has yet to be translated to *in vivo* implementations. It could provide an approach to increase the scalability of designed biological systems with information processing capabilities and to design robust, modular and scalable a biological computer as we propose herein.

We propose an implementation of an instruction memory in the form of a non-volatile DNA based memory. The addressing of this memory is performed with a program counter (PC), which is implemented as a synchronous biological counter. The proposed biological counter is composed of D flip-flop circuits in a master-slave configuration. This means that the counter can change its internal state only on the edge of the biological clock signal. The counter as PC is used to define the instruction memory location of the instruction that should be executed in the next processing step. The proposed circuit is thus able to execute biological programs sequentially in the same manner as *central processing unit* (CPU) or processor, which presents the core of each modern computer (see Figure 1).

**Figure 1:**
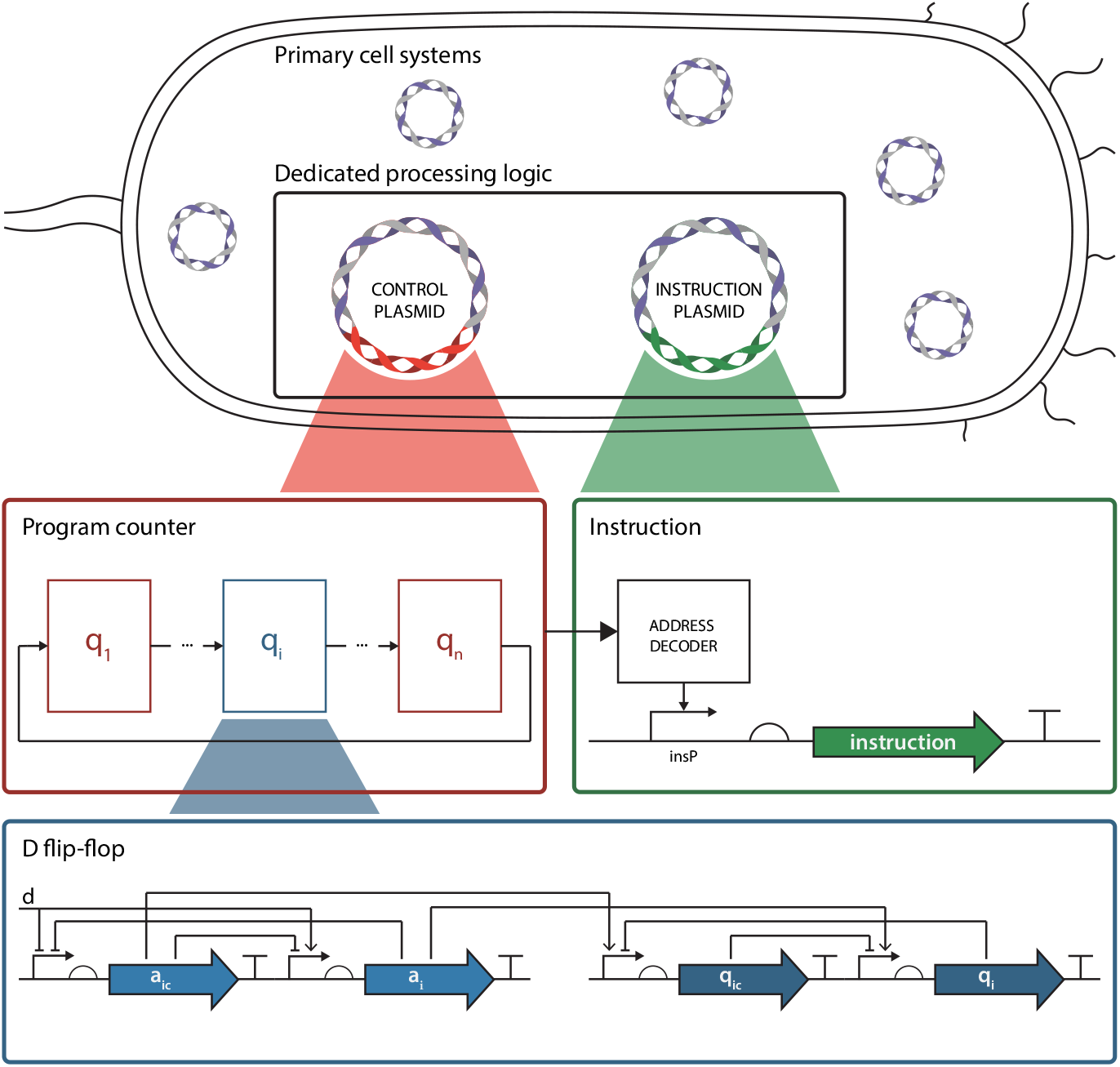
The proposed design of a biological processor in a biological host. The execution of instructions is governed by the state of the program counter implemented as a series of biological D flip-flops. A decoding logic is used to decode the current state of the program counter and to address and execute an instruction from the instruction memory.

## 2 Results

### 2.1 The clock signal can be generated by synthetic oscillators

All modern digital electronic systems are synchronized with a clock signal, which is an oscillatory signal with a predefined period. Synchronization is used to increase the predictability, scalability and speed of the response.

In biological systems, a clock can be generated by an oscillatory genetic circuit, such as a repressilator [9]. The majority of current synthetic biological information processing structures are implemented as asynchronous circuits [31]. This means that the switches within these circuits are not synchronized among each other. Genetic circuits generating oscillatory signals have already reached high reliability and reproducibility. For example, regular periodic switching was performed with recombinase-based molecular circuit [32]. Moreover, a lot of effort has been devoted to increase the reliability of existing synthetic biological oscillators [33] (see for example the extension of the repressilator described in [13]).

We propose a simulation framework that includes a clock generator of a sinusoidal signal, with constant amplitude, period and phase, which is consistent with the response of a repressilator. The proposed clock generator can be described with

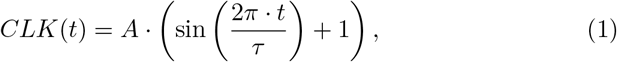

where *CLK*(*t*) is the clock signal, *t* is the time, *A* is the amplitude and *τ* the period of oscillations. Clock signal generator could alternatively be described as an ODE-based model of a repressilator or another biological circuit, which would not change the obtained results, but would increase the computational complexity of the simulation framework.

### 2.2 A synthetic oscillator synchronizes the counting events

Switching of the synchronous circuits should be limited to one switch per clock period, which is achieved with the synchronization of the response with the positive or negative edge of the clock signal. This allows us to synchronize the whole circuit with a single event and thus to increase the reliability of its response, which enables us to construct systems with larger complexity.

Several different versions of biological flip-flop have already been implemented, namely from the *toggle switch* [10] and *push-on push-off switch* [34] to the recently proposed *gated D latch* [35]. However, none of these structures have been synchronized with a clock signal.

We propose to use the Master-Slave D flip-flop sensitive to the positive edge of the clock signal as described in [29]. While the dynamics of asynchronous flip-flops is dependent solely on the *so-called* data inputs, their synchronous version also depend on the clock input, which synchronizes the flip-flop switches. Edge-triggered flip-flops coupled with biological oscillators can be further extended to biological counters as described in [29]. In digital circuits, a counter presents a canonical example of a sequential structure. Even though implementations of biological counters were already reported in the literature (e.g., see [36, 37]), these counters are still very limited in the sense of the number of events they are able to count/memorize. The main problem of existent implementations is their scalability and their sensitivity to the variability of data input signals. Data input pulse needs to be active long enough in order to perform the switch of the counter and at the same time short enough to prevent multiple switches [36]. It is hard or even impossible to generate a signal that precisely fits this demand within a noisy environment. On the other hand, we can overcome this limitation with a counter, which uses edge-triggered flip-flops changing their state only when (1) the data input signal is active and (2) when the clock signal changes either from inactive to active or vice versa.

A Master-Slave D flip-flop sensitive to the positive edge of the clock signal as described in [29] can be modelled with the following equations:

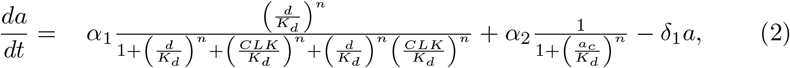

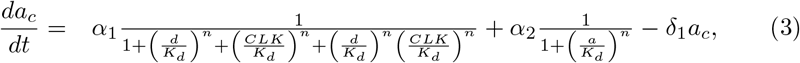

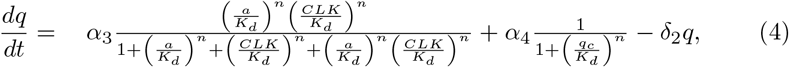

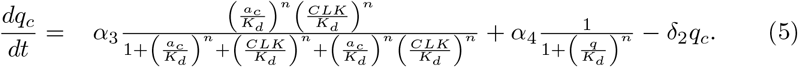

Here, *d* presents the data input and *CLK* the clock input. The parameters *α*_1_, *α*_2_, *α*_3_ and *α*_4_ are the expression rates of proteins with concentrations *a*, *a_c_*, *q* and *q_c_* respectively, *K_d_* is the dissociation constant, *n* is the Hill coefficient, and *δ*_1_, *δ*_2_ are the degradation rates of the observed proteins.

We can use the proposed biological flip-flop to design an arbitrary sequential biological circuit, such as a synchronous counter. A counter that counts up to 2^*m*^ events can be implemented using a sequence of *m* flip-flops. However, such an implementation requires a substantial amount of additional logic gates. A compromise between the circuit complexity and the size of the counting sequence can be made with the implementation of a Johnson counter as described in [29]. This counter does not require any additional logic besides D flip-flops and uses the inverted output of the last flip-flop in the sequence as an input to the first flip-flop in the sequence (see Figure 2). It is able to count up to 2*m* events, where *m* is the number of flip-flops used.

**Figure 2:**
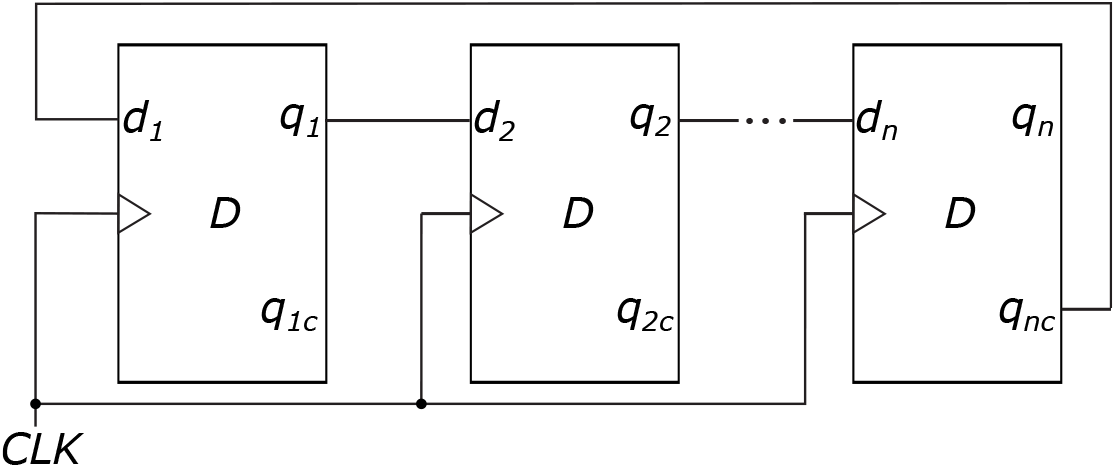
The implementation of a Johnson counter using D flip-flops. A John-son counter can count up to 2*m* events, where *m* is the number of flip-flops in the counter.

### 2.3 The Johnson counter as a program counter

In modern processors the instructions that need to be executed are stored within the *so-called* instruction memory. A program counter is a circuit, which enables us to sequentially execute the program stored in this memory. The counter should increase its value after a certain instruction has been executed, which can be interpreted as a direction to execute the next instruction in the memory.

Memory addressing is problematic in modern computer systems, since large decoders need to be implemented. For example, when decoding a memory with *n* address bits *n*/2^*n*^ decoders, i.e. decoders with *n* binary inputs and 2^*n*^ binary outputs, have to be used, which require the implementation of 2^*n*^ *n*-bit AND gates [28]. One of the benefits of the application of a Johnson counter as an addressing circuit is in the straightforward decoding of its current state using only two input AND gates independently of the size of the counter. Addressing of each memory location, however, still requires different two input AND gates. This means that *n* two input AND gates need to be implemented, where *n* is the number of memory addresses (see Table 1 for an example of decoding logic for 3-bit counter).

**Table 1:** An example of a decoding scheme used in a combination with a 3-bit Johnson counter. The counting sequence in this example is 000 → 100 → 110 → 111 → 011 → 001 → 000. Signals *q*_1_, *q*_2_ and *q*_3_ describe the state of the counter and serve as address inputs to the instruction memory, which identifies the location of the instruction to execute with the given decoding functions.

| $q_1$ | $q_2$ | $q_3$ | decoding function | instruction |
| --- | --- | --- | --- | --- |
| 0 | 0 | 0 | $\bar{q}_1$ AND $\bar{q}_3$ | $I_1$ |
| 1 | 0 | 0 | $q_1$ AND $\bar{q}_2$ | $I_2$ |
| 1 | 1 | 0 | $q_2$ AND $\bar{q}_3$ | $I_3$ |
| 1 | 1 | 1 | $q_1$ AND $q_3$ | $I_4$ |
| 0 | 1 | 1 | $\bar{q}_1$ AND $q_2$ | $I_5$ |
| 0 | 0 | 1 | $\bar{q}_2$ AND $q_3$ | $I_6$ |

The decoding function of the instruction *I_i_* can be modelled with the following equation

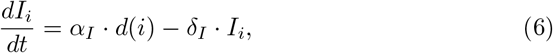

where parameters *α_I_* and *δ_I_* describe the production and degradation rates of the instructions and *d*(*i*) is the decoding function for the instruction *I_i_*, which has the following form

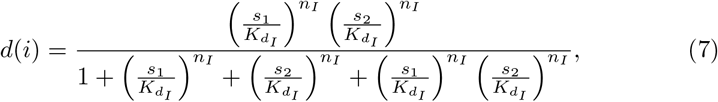

where 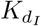 is the dissociation constant, *n_I_* is the Hill coefficient and *s*_1_ and *s*_2_ represent the concentrations of the inputs to the decoding function in accordance with Table 1.

### 2.4 The proposed circuits constitute a biological processor

State of the biological Johnson counter, i.e. program counter in this specific example, can be used as an address input to the instruction memory. The instruction memory is composed of instructions and of decoding functions, which are used to identify the instruction to execute (see Table 1). Decoding functions can be implemented through a promoter, which requires two inputs to be present at the same time (AND gate) as described with the model in Equation 6. When the promoter is induced it starts transcribing the DNA sequence that describes the instruction located at the specific location of the instruction memory. The program is thus *hardwired* in the DNA and the condition of execution of each specific instruction in this program is predefined with a specific value of the program (Johnson) counter. The circuit composed of the described subunits, namely the clock generator, the program counter and the instruction memory with the address decoding logic, constitutes a biological processor, which executes the instructions hardwired in the DNA program. The scheme of the proposed processor is presented in Figure 3.

**Figure 3:**
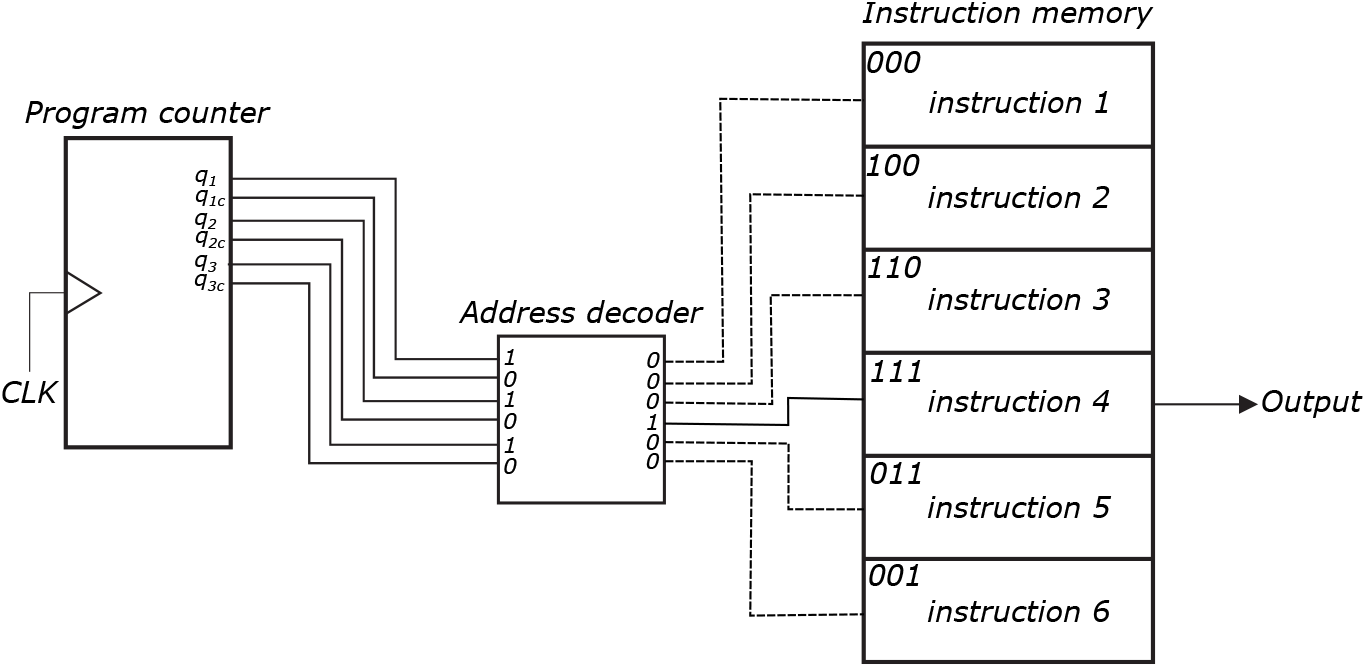
A scheme of the proposed biological processor. The program (Johnson) counter changes the state only on the positive edge of the clock signal (CLK) produced by the clock signal generator. The proposed processor uses address decoding logic to identify the location in the instruction memory and the instruction to execute in the given clock period.

### 2.5 The proposed biological processor is robust and scalable

We analysed the described topology using the methodology previously reported in [30]. This methodology has already been applied to the analysis of the biological D flip-flop [29], the repressilator [9] and the AC-DC circuit [38]. It can be used to assess the parameter space for which a topology reflects predefined dynamics described with a given fitness or cost function. Based on the parameter space assessment we can estimate the overall robustness and sensitivity of the topology to external perturbations. We performed the analysis on the ODE (ordinary differential equations) based models of the different biological processor topologies, each composed of a clock signal generator, a program counter and an instruction memory. We analysed versions composed of the 1-, 2-, 3- and 4-bit program counters, which can be used to address 2, 4, 6 and 8 instructions in the instruction memory, respectively. We were thus able to assess the robustness of these topologies as well as their scalability. For each, we assessed the feasible parameter space composed of parameter values for which the response is in accordance with a correct response.

We measured the correctness of the response with a cost function, which penalized the behaviour that deviates from the expected. We presumed that each address of the instruction memory contains an instruction for expression of a different protein. The cost function observed if these proteins were in fact expressed in a correct order. Moreover, we optimized the amplitude of expression of each instruction as well as of outputs for each of the flip-flops in the topology. The cost function *E*(*θ*) can be described as

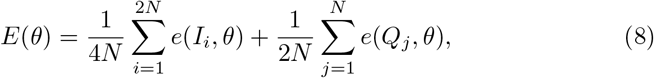

where *θ* defines the analysed parameter set, *N* the number of flip-flops in the topology, *I_i_* is the optimal response of the *i*-th instruction in the observed time-points, *Q_j_* the optimal response of the *j*-th flip-flop output in the observed timepoints, and where

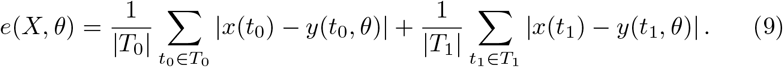

Here *X* describes the optimal course of the selected signal in the observed time-points (*X* = {*x*(*t_i_*)}), *T*_0_ includes all the timepoints, for which the condition *x*(*t*) ≤ 0.1 · max(*X*) is fulfilled, *T*_1_ includes all the timepoints, for which the condition *x*(*t*) > 0.9 · max(*X*) is fulfilled, and *y*(*t, θ*) presents the value of the signal in the given timepoint *t* obtained with the simulation on the parameter set *θ*.

We integrated the proposed cost function within the genetic algorithm (GA) optimization framework as described in [30]. We performed the parameter search on the 1-, 2-, 3- and 4-bit topologies. Solutions were interpreted as feasible if their cost functions did not exceed a certain threshold, which presents the average deviation of the simulation results from the optimal solution. The value of the cost function as well depends on the proportion of the switches that are performed during the simulation, since the deviance from the optimal trajectory is the largest during the switch. In order to obtain results that are directly comparable among different topologies, different threshold values were used for each of the topologies. The proportion of the switches between logic states is the largest in 1-bit topology. The most permissive threshold value was thus used here (i.e. 30 % of the maximal value of the optimal signal). On the other hand, the most stringent threshold value was applied to the 4-bit topology (i.e. 17 % of the maximal value of the optimal signal).

The optimization framework was able to produce feasible parameter spaces for each of the processor topologies, which we analyzed further. Figure 4 presents examples of viable solutions for 1-, 2- and 3-bit topologies. GAs sample the parameter values surrounding the optimal response of the system. In order to thoroughly sample the solutions which reflect the correct response of the system, local sampling around the optimal solutions is performed as described in [30]. The parameter ranges that reflected desired response in each topology are presented in Figure 5. Boxplots indicate that each of the parameters ranges for several orders of magnitude and thus indirectly indicate the robustness of the proposed design. Furthermore, we analysed the scalability of the obtained solutions. This can be analysed as the compatibility of the solutions obtained on the smaller topologies to the larger topologies. We assessed the compatibility among the solutions with the evaluation of cost functions across different topologies using parameter values obtained with the optimization of a specific topology. Results of this analysis are presented in Figure 6. Our results indicate that the solutions are mostly transferable among different topologies. However, solutions of 1-bit topology are only partially compatible with 2-, 3- and 4-bit topologies and vice versa. This indicates that the dynamics of the 1-bit topology is substantially different than other topologies. Solutions obtained with larger topologies, i.e. 3- and 4-bit topologies, are not always compatible with smaller topologies, i.e. 2- and 3-bit topologies. On the other hand, solutions obtained with the 2-bit topology are always applicable to 3- or 4-bit topologies and solutions obtained on the 3-bit topology are always applicable to the 4-bit topology. This shows that larger topologies reflect larger robustness and indicates the scalability of the proposed design.

**Figure 4:**
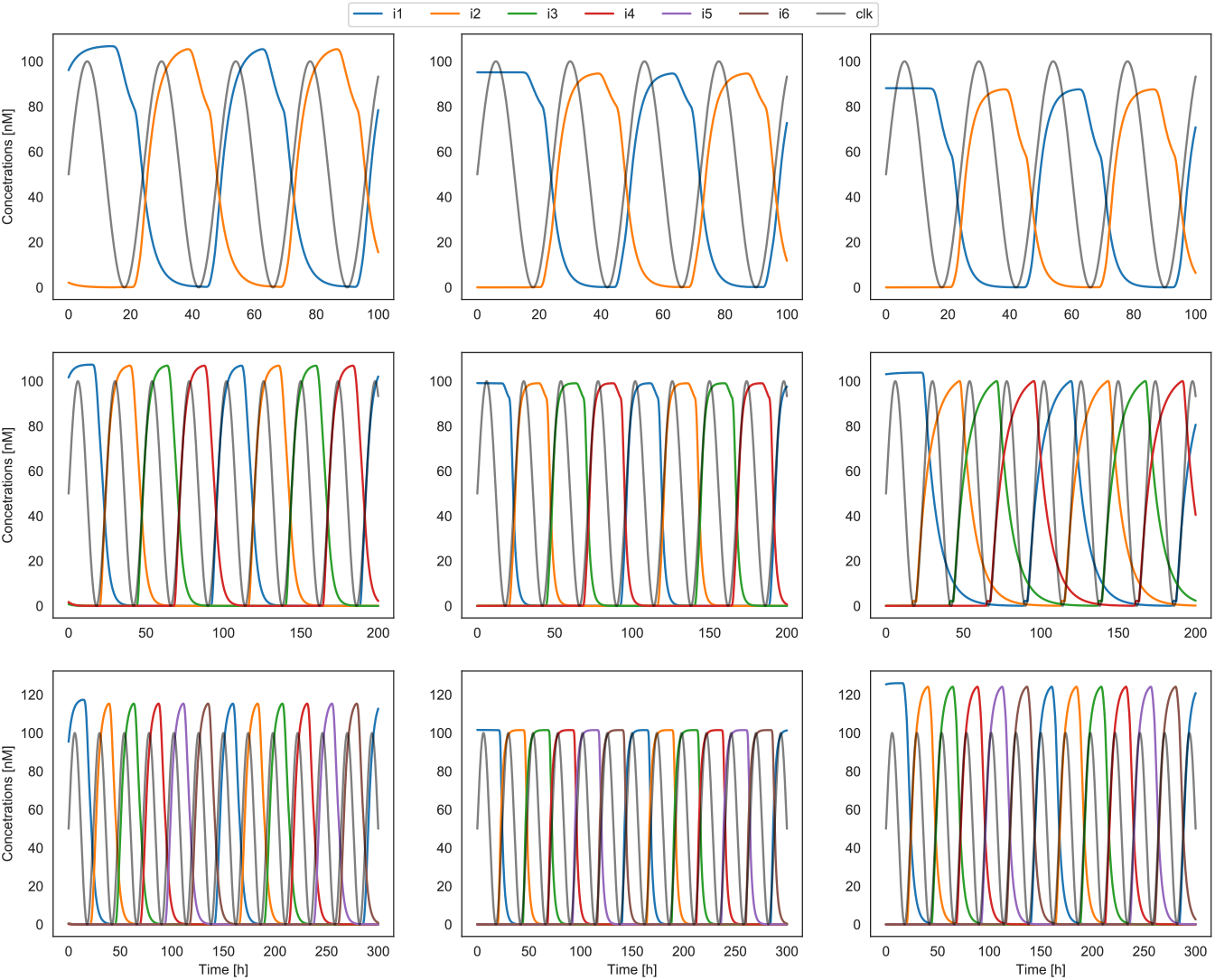
Examples of viable solutions for different topologies. The first, second and third row represent the simulation results of viable solutions for 1-, 2- and 3-bit topology, respectively. The signals *i*1 to *i*6 present the addressed instruction memory locations and *clk* presents the clock signal used for the synchronizationn.

**Figure 5:**
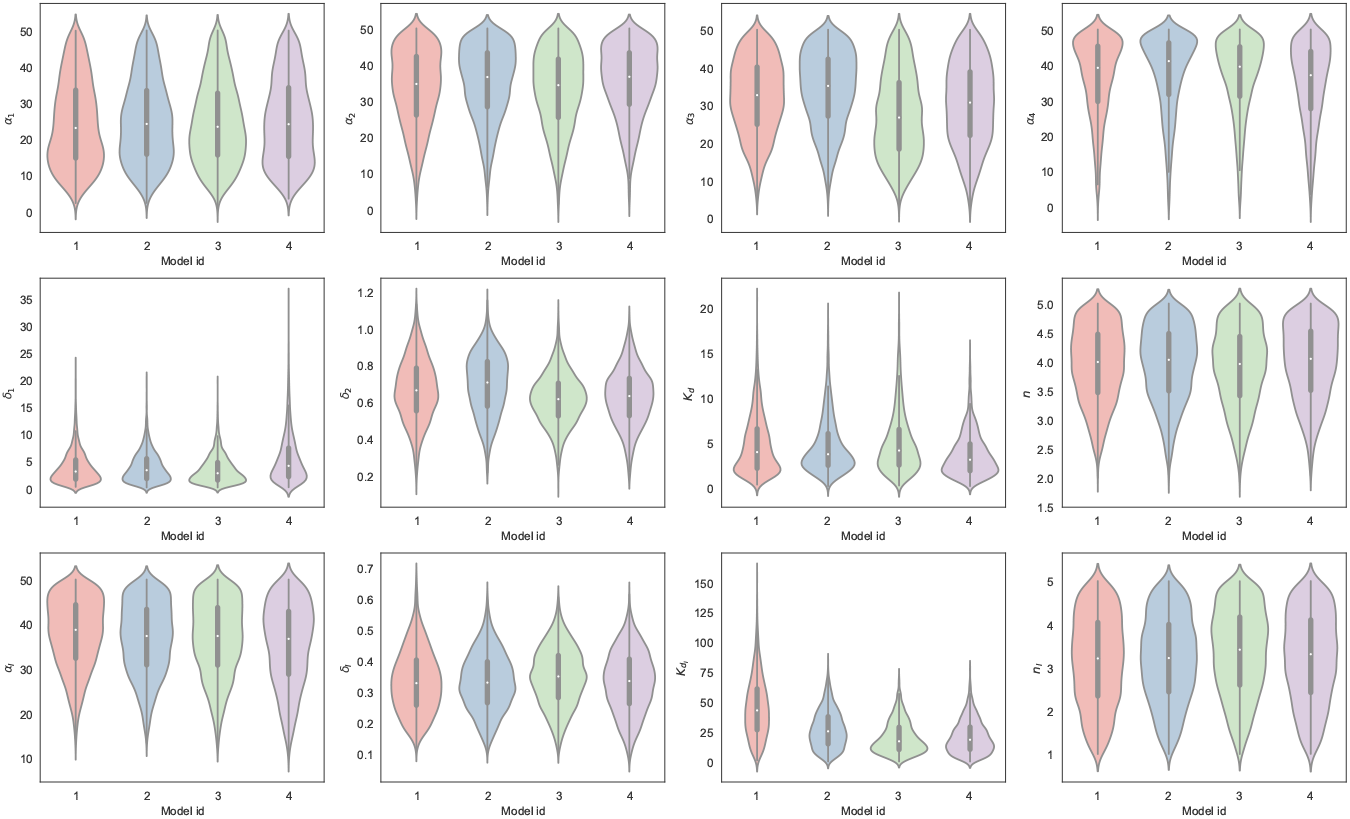
Parameter ranges for feasible solutions sampled around the optimal parameter values.

**Figure 6:**
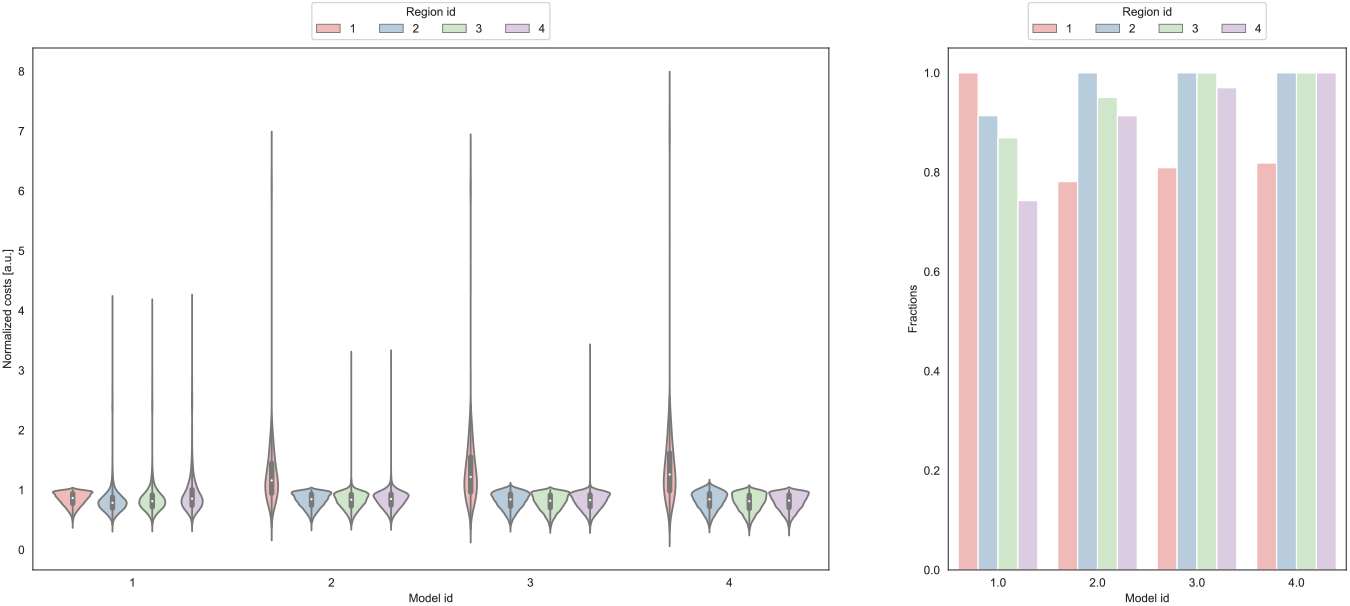
The assessment of the design scalability. We assessed the compatibility among the solutions with the evaluation of cost functions across different topologies (*Model id*) using parameter values obtained with the optimization of a specific topology (*Region id*). The figure on the left presents the distributions of values of normalized cost values across topologies. The figure on the right presents the fraction of feasible solutions obtained by using feasible parameter regions from a specific topology on different models. The cost values are normalized by the feasibility thresholds.

Another question that we wanted to investigate is how do the proposed topologies cope with the perturbations caused by the intrinsic noise, which is omitted in ODE-based deterministic simulations, but can be indirectly described with stochastic simulations using stochastic simulation algorithm (SSA) [39, 40] or its adaptations. We additionally validated the obtained results with a quasi steady-state approximation (QSSA) [41] of the SSA. This approximation enabled us to directly project the kinetic parameters assessed using the deterministic models to their stochastic equivalents. We selected points from the viable solution spaces and analysed their stochastic response. Points sampled from the feasible parameter regions pertained the expected behavior with similar characteristics also in the stochastic simulations. Figure 7 presents examples of the results of stochastic simulations performed with the same parameter values as used in Figure 4.

**Figure 7:**
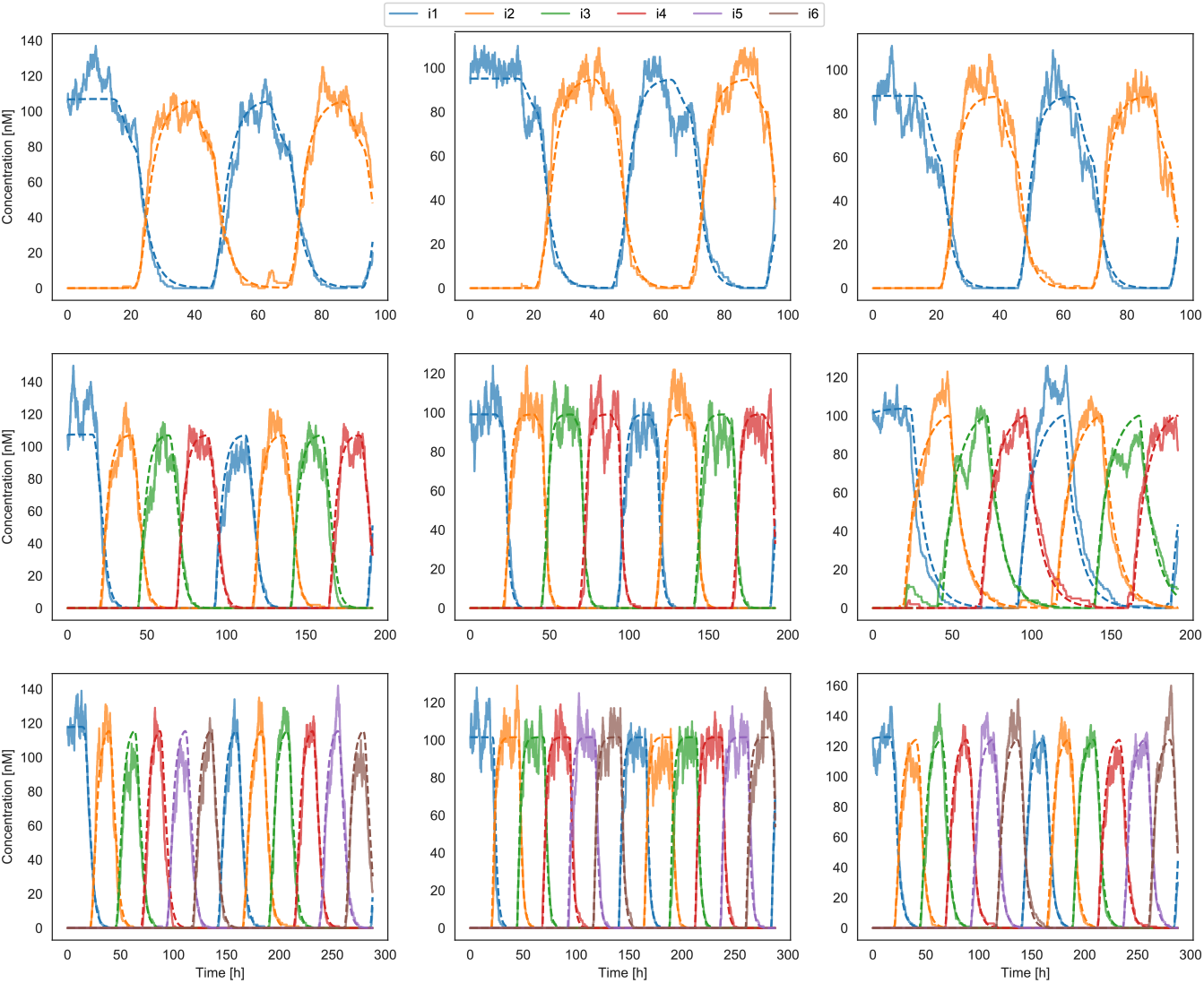
A comparison of the results obtained with deterministic and stochastic simulations. The first, second and third row represent the simulation results of viable solutions for 1-, 2- and 3-bit topology, respectively. Solid lines present the results of stochastic simulations and dashed lines present the results of deterministic simulations.

### 2.6 The biological processor can execute different programs

The proposed biological processor can execute different instructions, which compose the programs stored within the instruction memory. Some of these instructions, which as well present the basic instruction set of conventional processors, are described and discussed in the following sections.

#### 2.6.1 Basic instructions: generate and nop

The most basic instructions that are supported by the processor are `generate` and `nop` (no operation). While `generate` triggers the expression of a specific operand when the program counter points to its execution, `nop` performs nothing. It can be used as a placeholder instruction in the biological program and is usually applied when the cellular processor needs to wait for additional clock period until the next *real* instruction is executed.

#### 2.6.2 Addition

In the context of gene expression, addition can be interpreted as the expression of the same protein from two different genes, which are activated by the input operands. The concentration of expressed output operand is thus larger than in the case when the protein is expressed from a single gene. In the context of the proposed processor, addition, e.g. *C* = *A* + *B* or `add C,A,B`, can be implemented with an instruction, containing two genes, which both encode the result *C*. These are conditionally expressed, i.e. only in the presence of activator *A* for the first and only in the presence of activator *B* for the second gene.

#### 2.6.3 Subtraction

In the context of gene expression, result of the subtraction should be higher if only the first operand is present and lower if both or only the second operand is present. In the context of the proposed processor, subtraction, e.g. *C* = *A − B* or `sub C,A,B`, can be implemented with an instruction, containing a gene encoding protein *C*. This gene is expressed only if activator *A* is present and if repressor *B* is absent at the same time.

#### 2.6.4 Jumps

In order to allow jumps, which present the basis for implementation of *if-then sentence* and *loops*, we extended the proposed flip-flops with asynchronous inputs. These might be implemented with engineered proteases, which break down proteins presenting the *on* state of the flip-flop (i.e. *q* and *a*) in the case of asynchronous *set*, and break down proteins presenting the *off* state of the flip-flop (i.e. *not q* and *not a*) in the case of asynchronous *reset*. The ODE-based model of extended D flip-flop, can be described as

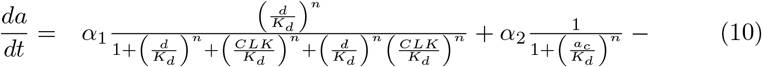

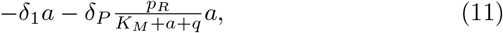

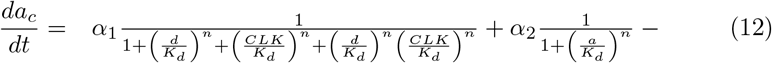

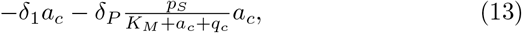

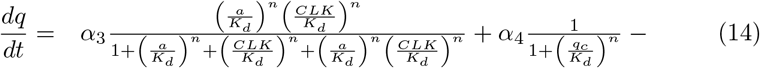

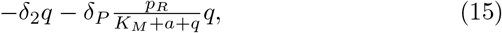

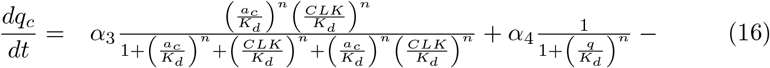

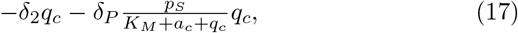

where *p_R_* and *p_S_* present the reset and set protease concentrations respectively, *δ_P_* is the rate of protease induced degradation, *K_M_* is its Michaelis constant and all other parameters correspond to those used in the Master-Slave flip-flop. Using this topology conditional jumps can be achieved with the expression of specific protease at the location in the memory from which jump should be performed. An example of the program executing an unconditional jump instruction is presented in Supplementary Figure 1.

Since the topology uses asynchronous set and reset inputs of proposed D flip-flop the jumps are performed asynchronously. i.e. without waiting for the next edge of the clock signal. Example in Supplementary Figure 1 shows that the instruction memory location from which the jump is performed is skipped and the processor jumps to the destination memory location immediately, i.e. within the same clock period.

Conditional jumps from a selected state can be achieved with the unconditional expression of a protease. However the activation of this protease is additionally regulated. Conditional jumps can thus be implemented using the proteases that are either activated by external inducer or inhibited by external inhibitor. Protease induction can be described with a Michaelis-Menten equation as

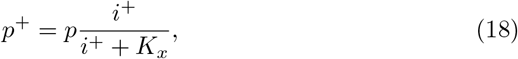

where *p* and *p*^+^ are the total and active protease concentrations respectively, *i*^+^ the inducer concentration and *K_x_* the dissociation constant. In a similar way we can describe the protease inhibition as

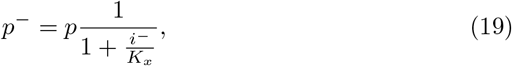

where *p*^−^ is the inactive protease concentration and *i*^−^ is the inhibitor concentration.

Simulation results in which the instruction at a specific memory location is skipped if the jump condition is fulfilled are presented in Supplementary Figure 2.

#### 2.6.5 Halt

The implemented jumps can be used to perform the so called `halt` instruction, which stops the processor at a given instruction memory location for an undefined amount of time. This can be implemented by performing an unconditional jump from the instruction memory location *i* to the location *i* − 1, i.e. to the previous location within the instruction memory. Supplementary Figure 3 presents the simulation results in which an unconditional jump is performed from a specific location in the biological program.

#### 2.6.6 If-then

If executes a certain sequence of instructions only when a condition is fulfilled. In this case a jump needs to be performed if the condition is not fulfilled. This can be implemented by using a condition as a protease inhibitor – the protease is active only if the inhibitor is absent and the jump is performed, i.e. the conditional instruction is skipped. Supplementary Figure 4 presents the simulation results in which a specific memory location is skipped if the condition is absent, i.e. if the condition is not fulfilled. Otherwise, i.e. if the condition is fulfilled, the instruction at this location is executed.

#### 2.6.7 While

A while loop executes a certain sequence of instructions as long as the condition is fulfilled. In order to implement a `while` loop two jumps need to be performed. Firstly, processor checks if the condition is fulfilled and in this case it starts executing the instruction within the loop. If the condition is not fulfilled, a jump to the memory location following the `while` loop is performed. This can be implemented in the same manner as the *if-then* sentence. However, an additional jump to the beginning of the while loop needs to be performed if the condition is still fulfilled. Supplementary Figure 5 presents the simulation results in which a specific memory location is addressed while the condition is fulfilled.

### 2.7 A biological compiler translates programs into a biological description

We implemented a compiler that automatically translates a sequence of given instructions into an ordinary differential equation based model. This can be simulated and different parameter values can be tested upon the model. The compiler is implemented in Python and accepts an arbitrary program composed of the instructions described in the preceding section. The program needs to be stored as a text file in which each line includes one or more instructions. Each line of the program is translated to a specific location in the memory, whereas first line is translated to the first address in the memory, second line to the second, etc. The compiler allows us to store an arbitrary number of instructions to the same memory location. These will be executed in parallel, i.e. within the same clock period. The programmer of the biological processor needs to keep in mind that hazards due to parallel execution might occur. The Python implementation of the compiler and interactive Python notebook examples with the user manual are available in Supplementary Data 1.

## 3 Discussion and conclusions

We described the computational implementation and analysis of a biological processor based on three biological information processing structures, namely a program (Johnson) counter implemented with Master-Slave D flip-flops, an instruction memory implemented with an instruction decode logic and a set of instructions that are expressed at each memory location, and a clock signal generator (see Figure 3). We have investigated the feasible regions of parameter values that yield expected behaviour for the proposed circuits. The obtained values comprise large solutions spaces in all analysed topologies and reflect the expected behaviour even when they are used in stochastic simulations. Moreover, our results show that the proposed design is scalable and could be used in more complex implementations of synthetic biological processing structures.

We demonstrated the possibilities to use the proposed processor to implement a set of different instructions, similar to the instructions available in the instruction sets of modern digital electronic processors. We have as well described the implementation of a compiler, which compiles program composed of a set of proposed instructions into an ODE-based model. The compiler can be used to construct more complex programs, which are automatically translated into computational models with simulation capabilities. These can be used to analyse the dynamics of each program more precisely and to fine tune the response of the system with a given set of parameter values. We believe that the proposed biological processor with the available instruction set and with the implemented compiler will find a vast range of application in modern age synthetic biology.

Our computational analyses indicate that the proposed implementation of a biological processor is feasible within the living organism. Current models describe the implementation of the processor within a single cell. However, this implementation could as well be performed in multiple cells coupled through quorum-sensing or other cell-cell communication mechanisms. Distribution of the processor between different cells synchronized with inter-cellular communication mechanisms would decrease the metabolic burden of heterologous proteins and their effect on cell growth [18]. Moreover, implementation in which the same functions would be distributed among different cells that would be synchronized through inter-cellular communication might increase the robustness of the system [24]. From the mathematical perspective the results that would be obtained with the so-called *population model*, would not significantly differ from the results described here. The computationally complexity of such extension would on the other hand drastically increase, which could present a problem with the efficient investigation of the whole possible parameter space.

Our future work will of course include several extensions of the proposed topology. In this work, we have applied the transcription based information processing capabilities of biological systems. Recently, a lot of focus has been devoted to the implementation of logic structures using engineered proteases, which could also be applied to the information processing in our context. These, however, should be analysed extensively in the proposed context. More complex applications of protease-based logic might prove to be problematic when designing more complex systems due to irreversibility of the proteolysis process.

In the proposed work we have analysed the synchronization of the biological processor with a synthetic clock generator. It might be feasible to couple the processor with oscillatory signals that are already present within the natural environment, such as the circadian clocks. These reflect large robustness, low variability and do not require additional engineered parts for their implementation. They could thus prove to be more efficient in the synchronization of the response of the cellular processor and their integration into the synthetic logic also presents one of our future goals.

## 4 Methods

### 4.1 Mathematical modelling of the proposed biological circuits

We performed our analyses using ordinary differential equation (ODE) based mathematical models. We presumed the mRNA concentrations in quasi-steady state since their dynamics is faster than the dynamics of the proteins [42]. The biological relevance of these models is significant only with accurate values of kinetic parameters. We investigated the feasible parameter space, for which the response of the observed biological structure is in accordance with our expectations, with the methodology described in Section 4.2 and in [30]. The optimization framework was parallelized in order to increase the efficiency of the computational analysis. Results obtained with this methodology were validated with the execution of stochastic simulations based on stochastic simulation algorithm (SSA) [39, 40] and its quasi steady-state approximation (QSSA) [41]. These are able to account for heterogeneity and the intrinsic noise of the biological systems [43], but are computationally more expensive than ODE-based simulations.

### 4.2 Computational analysis and optimisation framework

We applied the methodology described in [30] in order to efficiently investigated the parameter ranges for which the observed system exhibits the expected response. The methodology is able to identify the so-called *feasible* parameter space by performing the following steps:

1. Global estimation of viable parameter regions using the sampling of parameter values and evaluation of the response of the system at each value in a combination with genetic algorithms [44] and a predefined cost/fitness function.
2. Efficient local sampling using principal component analysis as described in [45].
3. Robustness analysis based on an estimation of the feasible parameter regions.

The possible ranges of parameter values for which the analysis was performed are described in Supplementary Table 1.

## Data availability

All the code together with the data and figures used in this paper is available at https://github.com/mmoskon/BioProc under the Creative Commons Attribution License.

## Acknowledgements

The research was partially supported by the scientific-research programme Pervasive Computing (P2-0359) financed by the Slovenian Research Agency in the years from 2013 to 2023 and by the basic research project CholesteROR in metabolic liver diseases (J1-9176) financed by the Slovenian Research Agency in the years from 2018 to 2021. L.M. was supported by young researcher scholarship provided by the Slovenian Research Agency. Results presented here are in the scope of the Ph.D. theses that are being prepared by Ž.P. and L.M.

## Author information

### Affiliations

Faculty of Computer and Information Science, University of Ljubljana, Slovenia

## Contributions

M.Mo. conceptualized the designs, implemented the models and the compiler. Ž.P. and M.Mo. designed and implemented the computational optimization and analysis framework. M.Mo., Ž.P. and L.M. executed the computational analysis and analyzed the results. M.Mo. wrote the manuscript. L.M., M.Mr. and N.Z. provided critical feedback and helped shape the research, analysis and manuscript. All authors read and approved the final manuscript.

## Corresponding author

Correspondence to Miha Moškon.

## Ethics declarations

### Competing interests

The authors declare that they have no competing interests.

## Notes

https://github.com/mmoskon/BioProc

