## Supplementary material for "A Computational Design of a Programmable Biological Processor": Supplmentary Information

### 1 Supplementary Tables

**Supplementary Table** 1: Possible ranges of parameter values. The ranges were obtained from [1, 2, 3]. We excluded some of the extreme cases from these ranges, such as extremely long-lived or unstable proteins. $V_R$ describes the reaction space volume and $N_A$ the Avogadro constant.

| Parameter | Span | Unit |
|---|---|---|
| Transcription | $10^{-2}$ – 50 | $nM\ h^{-1}$ |
| Translation | $10^{-2}$ – 50 | $h^{-1}$ |
| Protein production | $10^{-1}$ – 50 | $nM\ h^{-1}$ |
| Protein degradation | $10^{-3}$ – 50 | $h^{-1}$ |
| mRNA degradation | $10^{-1}$ – 100 | $h^{-1}$ |
| Dissociation constant | $10^{-2}$ – 250 | $nM$ |
| Hill coefficient | 1 – 5 | – |
| Reaction volume ($V_R \cdot N_A$) | 1 | $nM^{-1}$ |

### 2 Supplementary Figures

- Supplementary Figure 1 presents the simulation results in which instruction at the memory location *i4* is skipped with the application of unconditional jump. We can see that the instruction at the memory location *i4* is skipped and processor jumps to the memory location *i5* immediately, i.e. within the same clock period.

- Supplementary Figure 2 presents the simulation results in which the instruction at the memory location *i4* is skipped if the jump condition is fulfilled (the upper row). If the condition is not fulfilled, the instruction at the memory location *i4* is executed (the bottom row).

- Supplementary Figure 3 presents the simulation results in which an unconditional jump is performed from the location *i5* to the location *i4*, which results in halt at the location *i4*.

- Supplementary Figure 4 presents the simulation results in which the memory location $i4$ is skipped if the condition is absent, i.e. if the condition is not fulfilled. Otherwise, i.e. if the condition is fulfilled, the instruction at the location $i4$ is executed.

- Supplementary Figure 5 presents the simulation results in which memory location $i4$ is addressed while the condition is fulfilled.

**Supplementary Figure** 1: Simulation results of the program in which an unconditional jump is used to skip an instruction at the memory location *i4*. Implementation of the instruction counter using flip-flops with asynchronous set and reset inputs allows the execution of unconditional jumps. The same program was executed on three different randomly selected solutions from the feasible parameter region obtained with the methodology described in [4]. The proteolysis parameters were set to the following values: $K_M = 100\ nM$, $\delta_P = 250h^{-1}$.

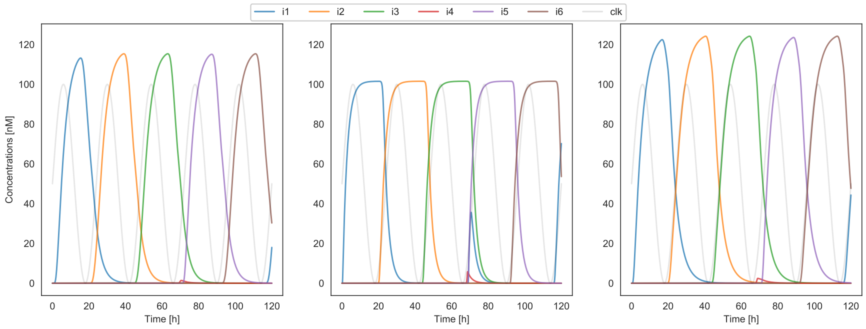

**Supplementary Figure** 2: Simulation results of the program in which a conditional jump is used to skip an instruction at the location *i4*. Implementation of the instruction counter using flip-flops with asynchronous set and reset inputs allows the execution of conditional jumps. The upper row presents the scenarios in which the jump condition is fulfilled (the protease inducer is present). The bottom row presents the scenarios in which the jump condition is not fulfilled (the protease inducer is absent). This behaviour can be interpreted as an *if-then* instruction. The program was executed on the same solutions as used in Supplementary Figure 1. The proteolysis parameters were set to the following values: $K_M = 100\ nM$, $\delta_P = 250h^{-1}$, $K_x = 0.1\ nM$.

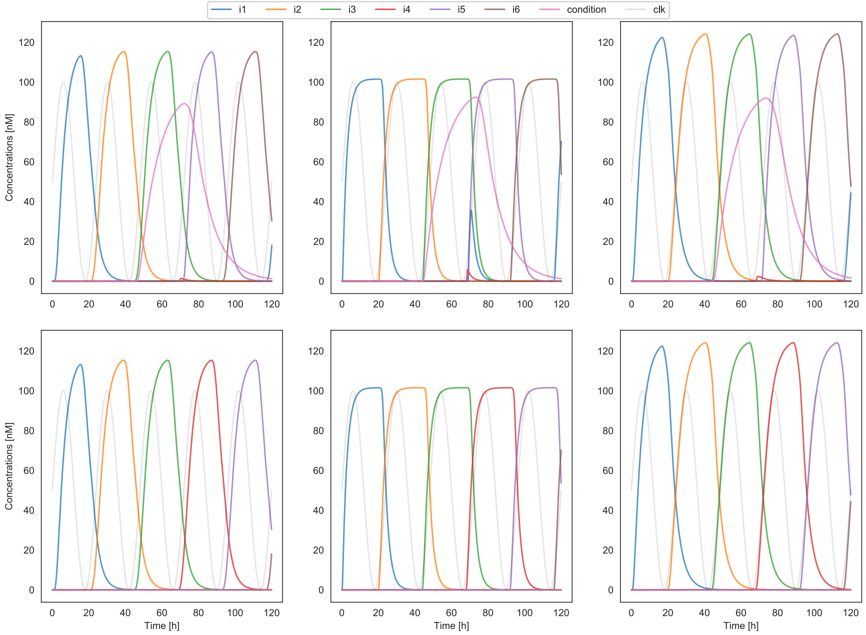

**Supplementary Figure** 3: Simulation results of the program in which a jump is used to halt the processor on the instruction at the memory location *i4*. The program was executed on the same solutions as used in Supplementary Figure 1. The proteolysis parameters were set to the following values: $K_M = 100\ nM$, $\delta_P = 250h^{-1}$, $K_x = 0.1\ nM$.

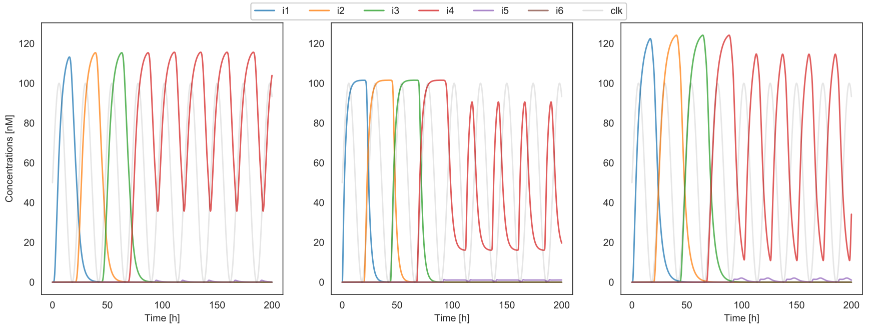


**Supplementary Figure** 4: Simulation results of the program in which instruction at the memory location *i4* is executed only if the condition is fulfilled (the upper row). The memory location *i4* is skipped if the condition not fulfilled (the bottom row). The program was executed on the same solutions as in Supplementary Figure 1. The proteolysis parameters were set to the following values: $K_M = 100\ nM$, $\delta_P = 250h^{-1}$, $K_x = 0.1\ nM$.

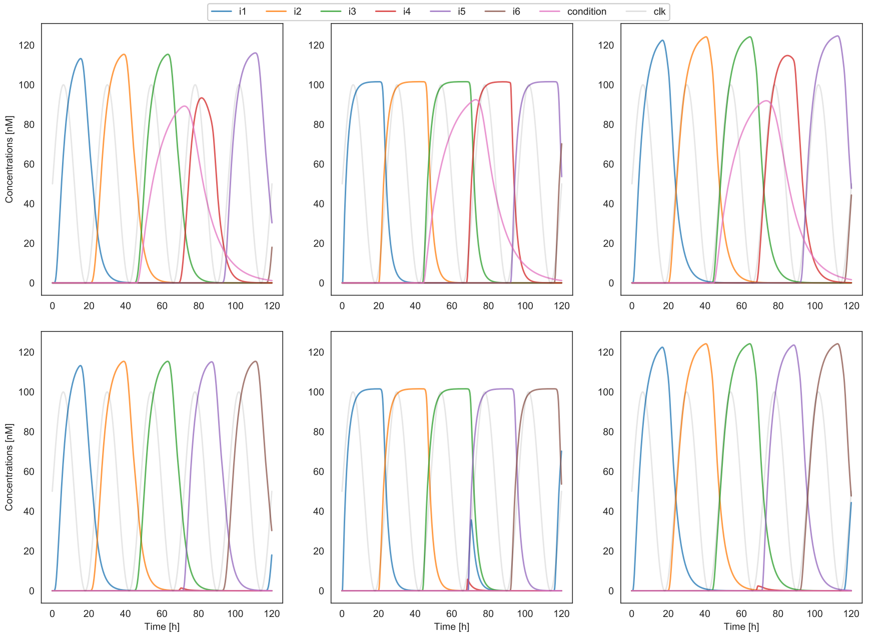

**Supplementary Figure** 5: Simulation results of the program in which operand $C$ is expressed as long as the condition is fulfilled. The program was executed on the same solutions as in Supplementary Figure 1. The proteolysis parameters were set to the following values: $K_M = 100\ nM$, $\delta_P = 250h^{-1}$, $K_x = 0.1\ nM$.

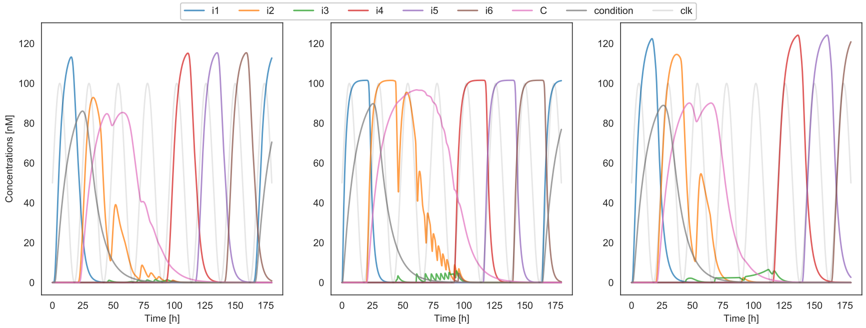

[2] Bionumbers. [Online]. Available: http://homepages.ulb.ac.be/dgonze/BIONUMBERS/bionumbers.html

[3] Bionumbersdatabase. [Online]. Available: https://bionumbers.hms.harvard.edu/search.aspx

[4] Ž. Pušnik, M. Mraz, N. Zimic, M. Moškon *et al.*, "Computational analysis of viable parameter regions in models of synthetic biological systems," *Journal of biological engineering*, vol. 13, no. 1, p. 75, 2019.
