## Supplementary Data for "A Computational Design of a Programmable Biological Processor"

### A Computational Design of a Programmable Biological Processor - Description of the biological compiler

The compiler accepts programs written as text files. Each line of a program is translated to a specific location in the memory, whereas first line is translated to the first address in the memory, second line to the second, etc. Multiple instructions can be given within the same line. In this case, instructions should be separated with semicolons ( `;` ).

Each program is compiled into an ordinary differential equation model with the simulation capabilities, which can be analysed further on.

#### How to use the biological compiler

The compiler is implemented in the Python module `generate_model.py` . You need to import the `generate_model` function from this module into a Python program.

In [1]:

```
from generate_model import generate_model
```

The `generate_model` function accepts the following arguments:

- `program_name` : name of the `txt` file in which the program is stored,
- `model_name` : name of the file in which the Python implementation of the model will be stored,
- `n_bits` : number of flip-flops in the Johnson counter that is used for the addressing of the instruction memory (defines the maximal size of the program),
- `prog_alpha` : maximal expression rate of proteins that are used as operands in the program,
- `prog_delta` : degradation rate of proteins that are used as operands in the program,
- `prog_n` : Hill coefficient defining the expression activation of proteins that are used as operands in the program,
- `prog_Kd` : dissociation constant defining the expression activation of proteins that are used as operands in the program.

An example of the function call is

In [2]:

```
n_bits = 4
generate_model("programs\\program_add.txt", "test_add", n_bits, 10, 0.1, 2, 10)
```

The upper call translates the program from the file `programs\program_add.txt` to the Python implementation of ordinary differential equation-based model named as `test_add.py` . This model can be imported using the `importlib` library.

In [3]:

```
import importlib
model = importlib.import_module("test_add")
```

The `generate_model` function also generates the list of operands that are used in the model. It stores this list in the file `model_name+"description.txt"` . You can read this file and display the operands used in the program.

In [4]:

```
f_description = open("test_add"+"description.txt")
ops = f_description.readline().strip().split(",")[:-1]
f_description.close()
ops
```

Out[4]:

```
['A', 'B', 'C']
```

In order to simulate the dynamics of the obtained model, you still need to define the remaining parameter values used for a simulation. You can obtain the most of the parameter values with the optimization framework proposed in *Pušnik et al., 2019*. Some feasible values are stored in the file `selected_points.txt`

In [5]:

```
import numpy as np
points = np.loadtxt('selected_points.txt')
params = points[0]
```

The first 8 parameters define the dynamics of flip-flops and the remaining parameters define the dynamics of the addressing logic.

In [6]:

```
params_ff = list(params[:8])
params_addr = list(params[8:])
```

You still need to define the proteolysis parameters used for asynchronous set and reset D flip-flop inputs and dissociation constant used in conditional jumps.

In [7]:

```
delta_P= 250
K_M = 100
K_X = 0.1

params_proteolysis = [delta_P, K_M]
params_condition = [K_X]
```

You can now simulate the dynamics of the program with the `odeint` function from the `scipy` module `integrate` .

In [8]:

```
from scipy.integrate import odeint
T = np.linspace(0, 100, 1000)
Y0 = np.array([0]*(n_bits*6+len(ops)))
Y = odeint(model.model, Y0, T, args= (params_ff + params_proteolysis + params_addr + params
```

Here `T` includes the timepoints in which results of a simulation will be sampled and `Y0` presents the initial state of the system.

The simulation results are now stored in the matrix `Y`. The operands present the last (in our case three) columns of the matrix. You can analyse the results of your program's execution by plotting the operands' concentrations through time.

In [9]:

```
import matplotlib.pyplot as plt
for i,op in enumerate(ops):
    plt.plot(T,Y[:,-(i+1)], label = op)
plt.legend()
plt.show()
```

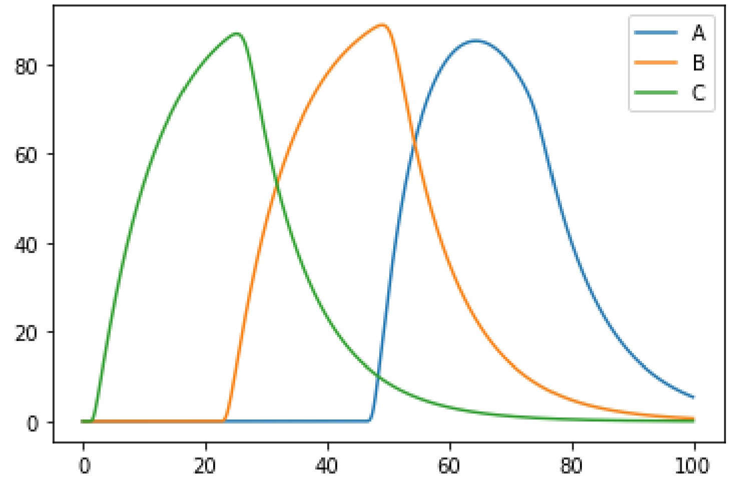


A faster alternative to test your programs is to use the `simulate_program` function from the `simulate_program` module, which automatically generates the program, runs the simulation and displays the simulation results. First you need to import this function into your module.

In [10]:

```
from simulate_program import simulate_program
```

The `simulate_program` function accepts the following mandatory arguments:

- `program_name` : name of the `txt` file in which the program is stored,
- `t_end` : simulation duration (in hours),
- `N` : number of samples,
- `params_ff` : flip-flop parameters,
- `params_addr` : addressing parameters,
- `params_proteolysis` : proteolysis parameters,
- `params_condition` : params used in conditional jumps,
- `params_prog` : params used for the expression of operands,
- `n_bits` : number of flip-flops in the Johnson counter that is used for the addressing of the instruction memory (defines the maximal size of the program).

An example of the function call and its output is

In [11]:

```
params_prog = [10, 0.1, 2, 10]
simulate_program("programs\\program_add.txt",
                 100,
                 200,
                 params_ff,
                 params_addr,
                 params_proteolysis,
                 params_condition,
                 params_prog,
                 4)
```

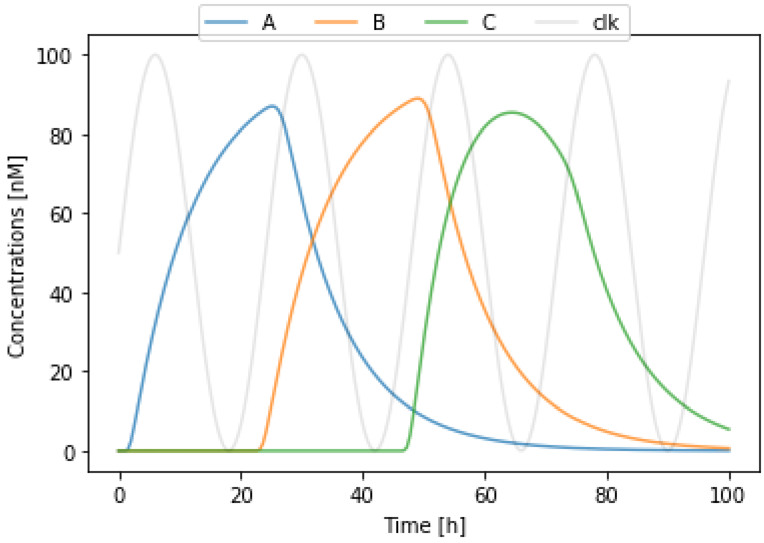


### Currently supported instruction set

##### `nop`

Only used as a placeholder instruction.

##### `generate op1`

Expression of operand with the name `op1` is triggered.

##### `add op1, op2, op3`

Performs the addition of the operands `op2` and `op3` and stores the result to `op1` :

`op1<-op2+op3`

##### `sub op1, op2, op3`

Performs the subtraction of the operands `op2` and `op3` and stores the result to `op1` :

`op1<-op2-op3`

##### `if condition instruction`

Executes the `instruction` if the concentration of operand `condition` is high. Can be followed by multiple instructions separated by semicolons ( `;` ), e.g., `if D generate A; generate B; add C,A,B` .

##### `do-while condition instruction`

Instruction `instruction` is always executed in the first loop transition and is executed as long as the operand `condition` is high.

##### `while condition instruction`

Instruction `instruction` is executed only if and as long as operand `condition` is high.

##### `halt`

Halts the processor at current instruction memory location.

### Examples

The following examples present the application of the proposed compiler on simple biological programs. The compiler is used to translate these programs into ordinary differential equation models, which are than used to simulate the dynamics of the program in dependence on the given set of kinetic parameters.

#### Initialization

##### Imports

In [12]:

```
from simulate_program import simulate_program
import numpy as np
import seaborn as sns

sns.set_style("white")
```

##### Simulation parameters

In [13]:

```
t_end = 160 # duration in hours
N = 1000 # number of samples to display
```

##### Program parameters

In [14]:

```
n_bits = 4 # number of bits in the instruction memory

# parameters defining the expression of operands
prog_alpha = 10
prog_delta = 0.1#0.01
prog_n = 2
prog_Kd = 10
params_prog  = prog_alpha, prog_delta, prog_n, prog_Kd

# proteolysis and induction of protease (conditional jumps)
delta_P= 250
K_M = 100
K_X = 0.1
```

##### Processor parameters

Use the parameter values that were obtained with the optimization framework proposed in *Pušnik et al., 2019*.

In [15]:

```
points = np.loadtxt('selected_points.txt')
params = points[0]

params_ff = list(params[:8])
params_addr = list(params[8:])

params_proteolysis = [delta_P, K_M]
params_condition = [K_X]
```

#### Examples of simple programs

Few examples are given to show how to compile a program stored in a text file and produce a model that can be simulated with the parameters defined above.

##### Addition

```
generate A
generate B
add C, A, B
```

In [16]:

```
program_name = "programs\\program_add.txt"
simulate_program(program_name, t_end, N, params_ff, params_addr, params_proteolysis, params
```

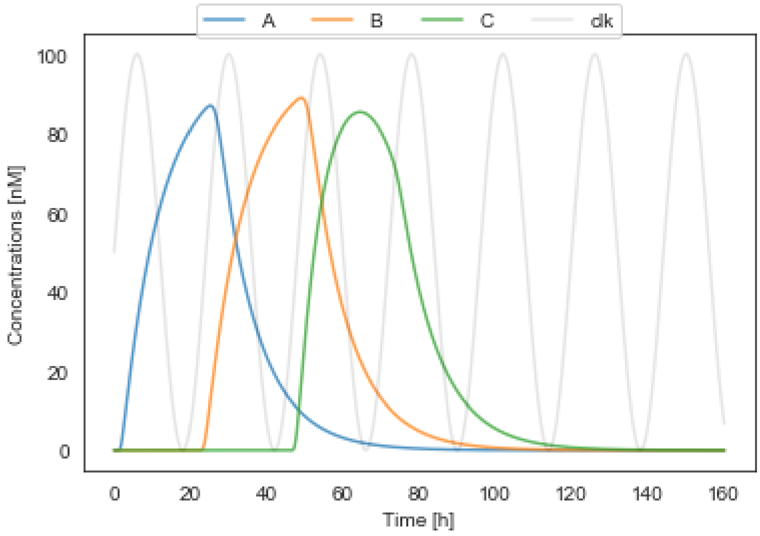


#### Subtraction

```
generate A
generate B
sub C, A, B
```

In [17]:

```
program_name = "programs\\program_sub.txt"
simulate_program(program_name, t_end, N, params_ff, params_addr, params_proteolysis, params
```

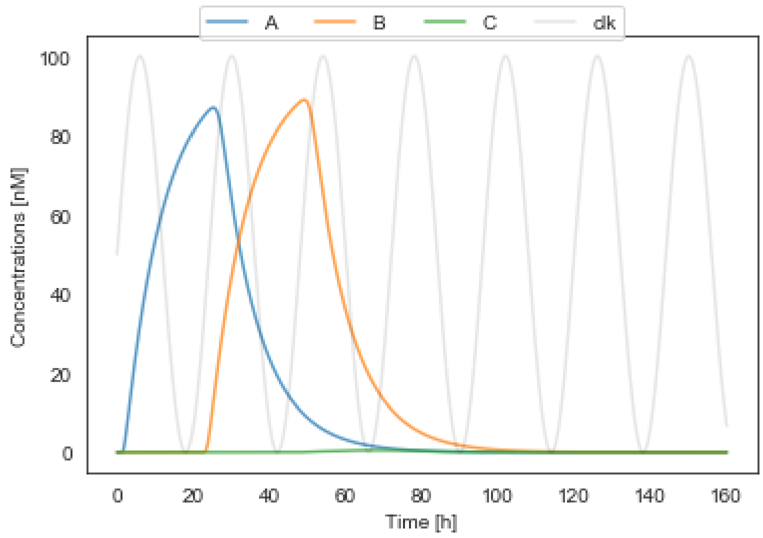


#### Subtraction 2

```
generate A
sub C, A, B
```

In [18]:

```
program_name = "programs\\program_sub_one.txt"
simulate_program(program_name, t_end, N, params_ff, params_addr, params_proteolysis, params
```

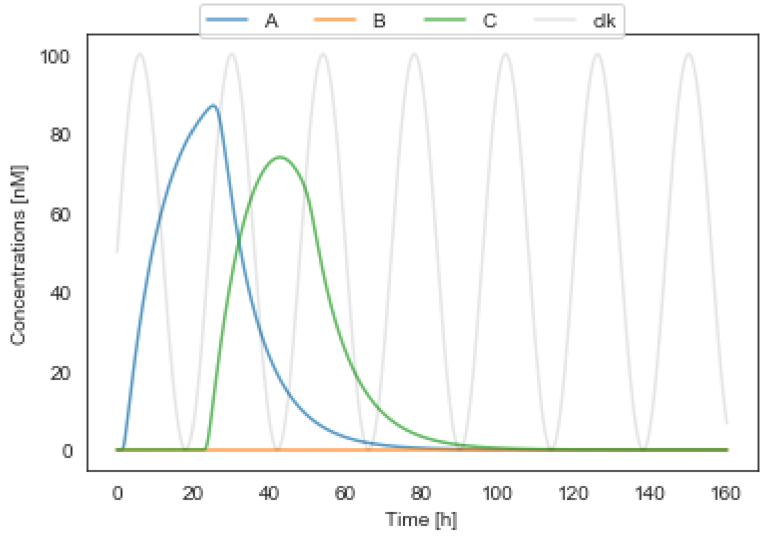


#### Addition and halt

```
generate A
generate B
add C, A, B
halt
```

In [19]:

```
program_name = "programs\\program_add_halt.txt"
simulate_program(program_name, t_end, N, params_ff, params_addr, params_proteolysis, params
```

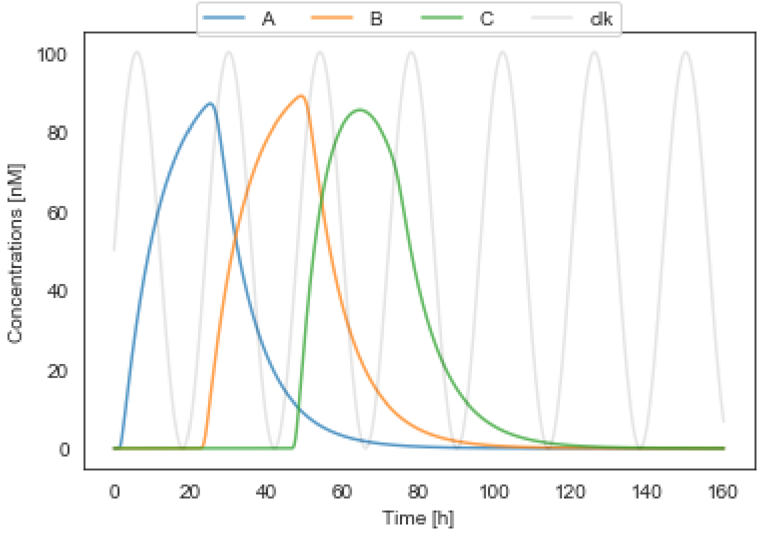


#### Multiple additions, subtraction and halt

This example demonstrates the execution of multiple instruction in the same clock period.

```
generate A; generate B; add E, A, B
add C, A, B; sub D, A, B
halt
```

In [20]:

```
program_name = "programs\\program_add_multi.txt"
simulate_program(program_name, t_end, N, params_ff, params_addr, params_proteolysis, params
```

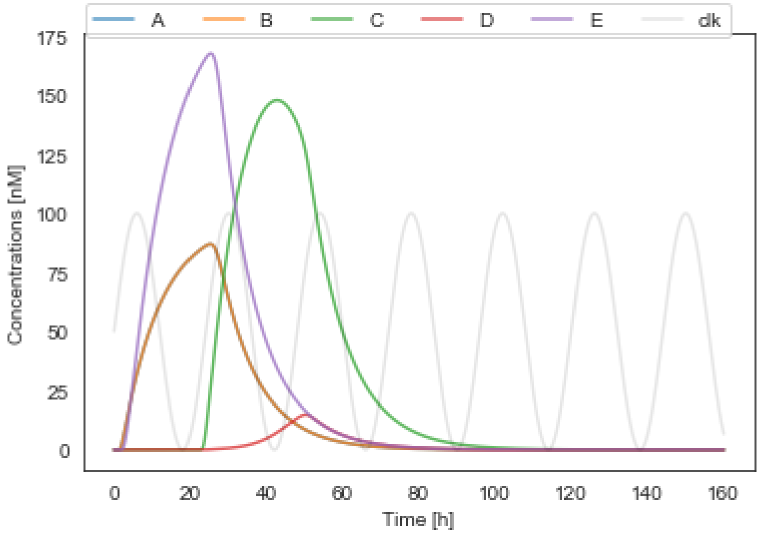


#### If-then

Condition is fulfilled

```
generate A
generate B
if B add C, A, B
```

In [21]:

```
program_name = "programs\\program_if.txt"
simulate_program(program_name, t_end, N, params_ff, params_addr, params_proteolysis, params
```

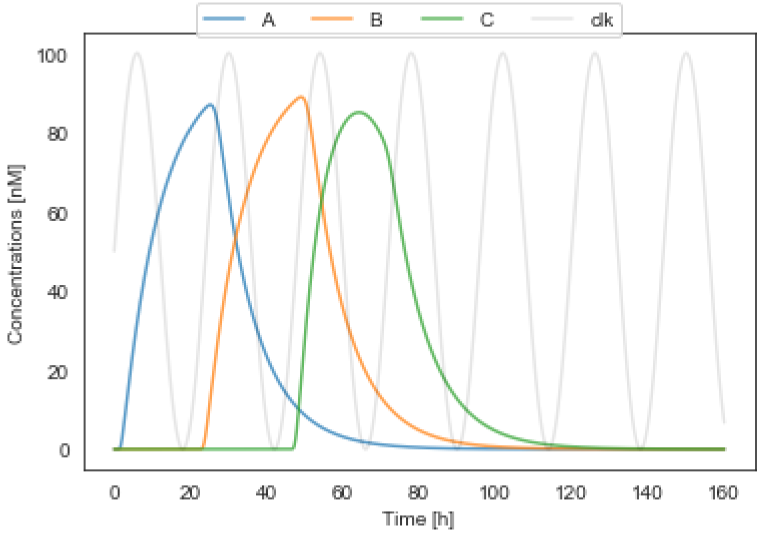


Condition is not fulfilled

```
generate A
if B add C, A, B
```

In [22]:

```
program_name = "programs\\program_if_false.txt"
simulate_program(program_name, t_end, N, params_ff, params_addr, params_proteolysis, params
```

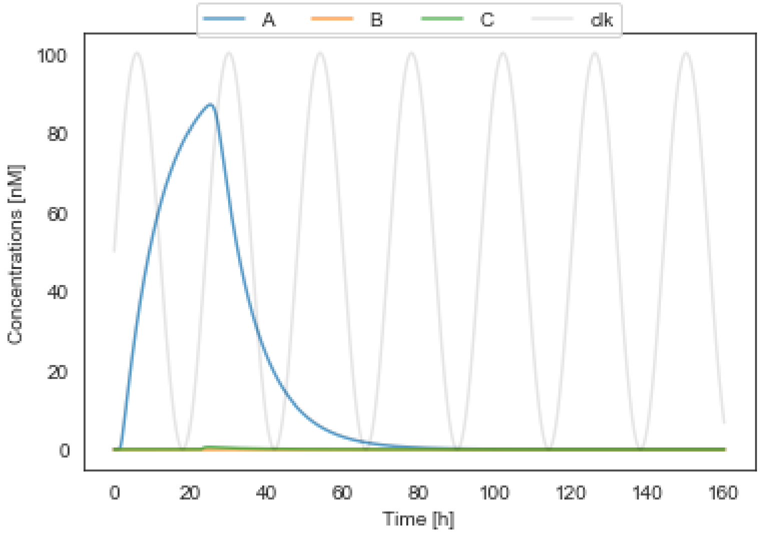


#### While

```
generate A
while A generate C
```

In [23]:

```
program_name = "programs\\program_while.txt"
simulate_program(program_name, t_end, N, params_ff, params_addr, params_proteolysis, params
```

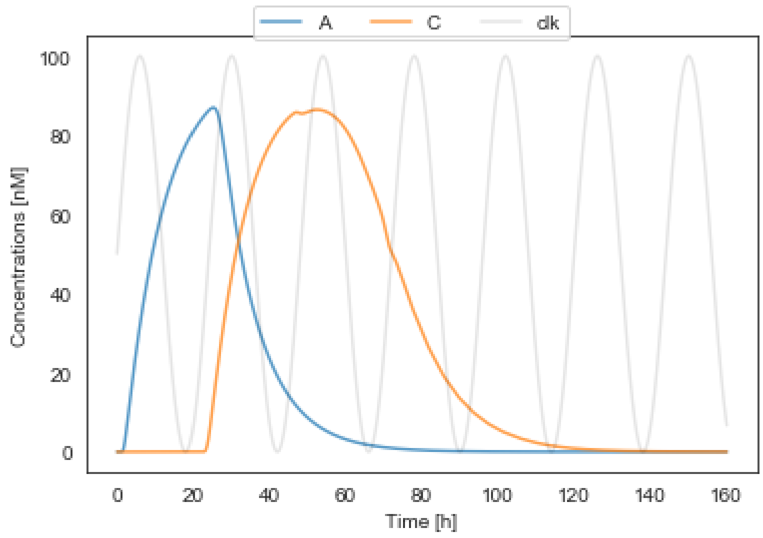
